# Diffsig: Associating Risk Factors With Mutational Signatures

**DOI:** 10.1101/2023.02.09.527740

**Authors:** Ji-Eun Park, Markia A. Smith, Sarah C. Van Alsten, Andrea Walens, Di Wu, Katherine A. Hoadley, Melissa A. Troester, Michael I. Love

**Affiliations:** UNC-Chapel Hill

## Abstract

Somatic mutational signatures elucidate molecular vulnerabilities to therapy and therefore detecting signatures and classifying tumors with respect to signatures has clinical value. However, identifying the etiology of the mutational signatures remains a statistical challenge, with both small sample sizes and high variability in classification algorithms posing barriers. As a result, few signatures have been strongly linked to particular risk factors. Here we present *Diffsig*, a model and R package for estimating the association of risk factors with mutational signatures, suggesting etiologies for the pre-defined mutational signatures. *Diffsig* is a Bayesian Dirichlet-multinomial hierarchical model that allows testing of any type of risk factor while taking into account the uncertainty associated with samples with a low number of observations. In simulation, we found that our method can accurately estimate risk factor-mutational signal associations. We applied *Diffsig* to breast cancer data to assess relationships between five established breast-relevant mutational signatures and etiologic variables, confirming known mechanisms of cancer development. *Diffsig* is implemented as an R package available at: https://github.com/jennprk/diffsig.

## Introduction

Somatic mutations are changes in the DNA sequence that occur throughout the life of an organism due to mutagenic processes such as DNA replication error, ultraviolet exposure, or smoking. Each mutagenic process is believed to leave a unique pattern in the DNA, the collective action of which creates “mutational signatures”^1^. Mutational signatures were initially introduced by Nik-Zainal *et al*. (2012) where breast cancer-specific signatures were discovered based on whole-genome sequenced (WGS) data using non-negative matrix factorization (NMF)^2^. A following large-scale study on 30 cancer types led to the construction of a compendium of detected signatures, the Catalogue Of Somatic Mutation In Cancer (COSMIC)^3^. Single base substitution (SBS) signatures from COSMIC are composed of 96 mutational contexts where each context represents a mutated base (C>A, C>G, C>T, T>A, T>C, and T>G) along with its immediate 5’ and 3’ bases (A, C, G, T). Each signature has different compositions of the mutational contexts, e.g. SBS2 is 98% composed of T[C>T]N contexts while SBS3 has a more uniform distribution of mutational contexts. Such mutational signatures are increasingly used to predict response to therapy, but also have an important role in understanding cancer development as they allow us to infer the mutational processes that occur throughout life and which sometimes lead to carcinogenesis^3^.

Numerous methods that allow for finding *de novo* signatures have been introduced. These methods detect mutational signatures based on various algorithms including Bayesian NMF ^4 5^, other variations of NMF ^6 7 9 10 11^, expectation-maximization^12^, or mixed-membership models^13 14^. These methods can detect novel signatures including ones that resemble the composition of mutational contexts in COSMIC signatures. Along with the discovered latent signatures, the estimated “contribution” is generated for each sample, which estimates how much each signature constitutes a sample’s count of somatic mutations.

Although many methods were successful in finding *de novo* signatures, identifying the etiology is not trivial. In previous studies, etiologies of the signatures were determined based on previous knowledge of somatic and germline mutations or cell lines^2 15^. Some studies also used statistical tests to detect the association between a risk factor and a signature, for example, Wilcoxon rank-sum tests comparing samples across binary variables, e.g. gender, and tobacco/alcohol usage^16^. Such comparison tests are usually performed directly on the number of mutation counts or the estimated contributions.

While such statistical methods presented a way to aid the process of discovering the associated mutagenic processes, a recent study by Yang et al. argued that such comparison tests may struggle with low power and efficiency due to the tests ignoring the variance of the estimated contributions^14^. Especially, when the total number of observed mutations is low, which may result from low sequencing coverage, low tumor purity, and/or low mutational burden, the accuracy of the estimated contributions may vary across samples and across signatures. To consider this, Yang et al. developed *HiLDA, which* utilizes a unified hierarchical latent Dirichlet allocation model within a Bayesian framework to identify *de novo* signatures and test the difference in contributions between two groups. This approach increases power as it directly models the latent contribution of signatures within each sample, though it only allows for testing associations between signatures and covariates in two group settings. Furthermore, it focuses on *de novo* signatures instead of leveraging pre-established signatures.

We propose *Diffsig*, a Bayesian Dirichlet-multinomial hierarchical model to estimate and test the association of risk factors and mutational signatures. While traditionally, one would consider risk factors that relate to a donor’s exposures, e.g. sunlight and tobacco usage, here we use the term “risk factor” generally to refer to any measured per-sample covariate that we wish to associate with mutational signatures. As with *HiLDA*, our proposed model is able to account for the heterogeneous uncertainty inherent with cancer patient cohorts where samples may have different numbers of observed somatic mutations due to varying sequence coverage and mutation burden. *Diffsig* can assess a large spectrum of risk factors including continuous and categorical risk factors. Moreover, multiple risk factors can have associations estimated simultaneously, which allows for unbiased estimation in the case of confounding. Unlike many other methods, we focus on finding the association between risk factors and the latent contribution of mutational signatures to each sample’s set of somatic mutations, hence we omitted the process of detecting *de novo* signatures and instead rely on any set of pre-defined mutation signatures that have reproducibly been relevant for breast cancer. Our model was evaluated with simulated datasets based on liver and breast cancer-related COSMIC signatures found from previous studies^17^ and with a breast cancer dataset generated by The Cancer Genome Atlas (TCGA) Research Network. We find our model can accurately capture biologically meaningful associations between risk factors and mutational signatures.

## Materials and Methods

### Bayesian Dirichlet-Multinomial Hierarchical Model

The goal of *Diffsig* is to estimate the associations between one or more risk factors and a predefined set of mutational signatures. In order to estimate the associations while accounting for sampling variance on heterogeneous samples, we use a Bayesian hierarchical Dirichlet-Multinomial model. As in the *HiLDA* model, we make use of a Dirichlet-Multinomial distribution; however, we do not include de novo signature detection as part of *Diffsig*, but instead only use pre-defined sets of signatures, which should be specified by the analyst based on literature or unsupervised analysis of the dataset. While a multinomial distribution can be used to model count data, the Dirichlet-Multinomial distribution, allows for potential differences in the latent contribution of mutational signatures for samples with the same risk factors.

The *Diffsig* model is built based on two main assumptions: 1) a sample’s mutational counts are composed of a specified set of mutational signatures, and 2) risk factors and mutational signatures have an association that can be approximated with a model that is linear in coefficients on the normalized exponential (softmax) scale. The hierarchical model is based on these assumptions (Figure 1) and our goal is to accurately estimate the association between risk factors and mutational signatures.

**Figure1.**
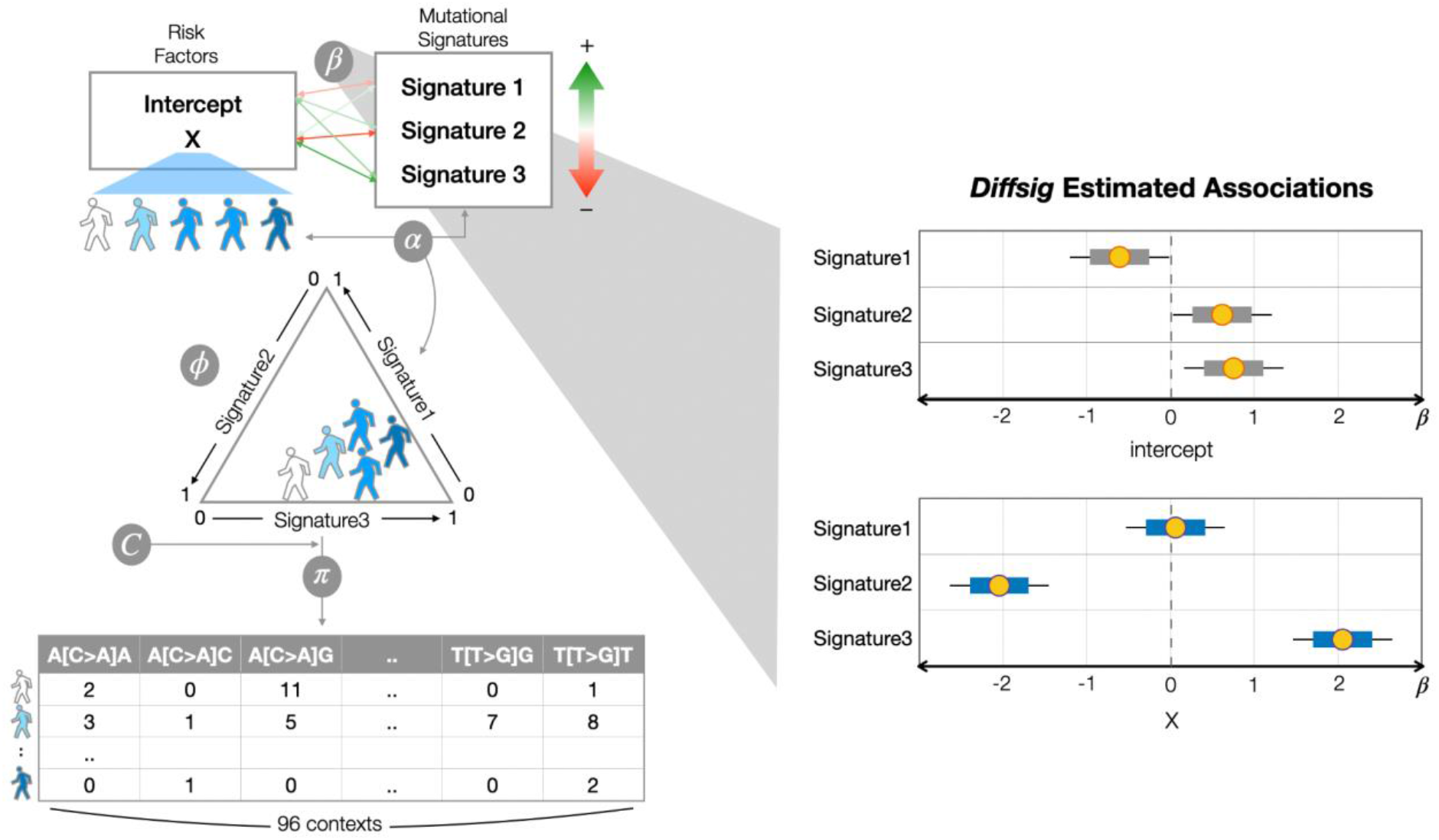
(Left Panel) Conceptual diagram of *Diffsig*, for detecting associations between risk factors and mutational signatures (with these target parameters defined as *β*). Each sample’s observed somatic mutations are composed of the association (*β*) and the sample’s risk factor traits (*X*). Associations between samples and signatures are defined as *α*. Even for samples with the same risk factor values, there exists variance among their signature distribution, here the center of a distribution per sample is diagrammed. The signature contribution is then multiplied by known transition probabilities for a set of reference signatures (*C*). Finally the resulting mutation probability *π* underlies the observed mutation counts (*Y*). (Right Panel) *Diffsig* estimates the associations *β* (yellow dots: point estimates, colored bars: 80% credible intervals, black lines: 95% credible intervals) for the left example.

For our model, three types of data are required: observed risk factors *X*, mutation counts *Y*_*N*×96_, and the pre-defined set of mutational signatures *C*_96×*K*_. The mutational signatures matrix contains columns representing the conditional probabilities of observing each type of mutational transition conditioning on signature, such that these probabilities add up to 1 for each signature *MS*_*k*,_ *k* ∈ 1, …, *K* where *K* is the number of signatures of interest. The observed risk factor data *X*_*N*×*M*_ should be complete for all *N* samples and *M* risk factors (*M* ≥ 2) including an intercept. In addition to the observed risk factors, we add an intercept that allows us to capture the baseline presence of mutational signatures 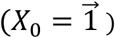. Risk factors can be in any form: binary, continuous, or categorical. For preprocessing, continuous variables were centered and scaled to have unit variance, and categorical variables with *p* levels were encoded as (*p*-1) dummy variables. With the given datasets, the association between risk factors and mutational signatures is captured with *β*, where *β*_*m*,._ is modeled with a normal distribution of mean 0 and standard deviation of *σ*_*β*_. The linear combination of risk factors *X* and the association coefficient matrix *β* then yields an association between samples and mutational signatures *α*_*i,j*_ where *n* ∈ 1, …, *N* and *k* ∈ 1, …, *K*.

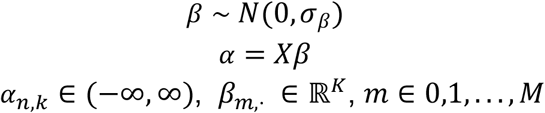

*α*_*N*×*K*_ therefore provides the relatedness of the samples and signatures, but is not in an easily interpretable space. Hence, *α*_*n*,._ is mapped with a normalized exponential function, i.e. softmax, for each row *n*, such that the values are compositional within each sample. Then a latent contribution for each sample *ϕ*_*n*,._ is modeled from a Dirichlet distribution on the softmax-ed *α* with a precision parameter *τ* that controls the concentration of the Dirichlet distribution around a vector of contributions *α*_*n*,._. The latent contribution *ϕ*_*K*×*N*_ allows us to incorporate the potential differences in the latent contribution for samples with the same risk factors. The latent composition of the samples for each signature *ϕ*_*k*,._ is then projected to the 96-dimensional mutational contexts *C* by taking the product of *ϕ* and the signature matrix *C*, giving the mutation probability *π*_*L*×*N*_. Finally, we assume the observed counts *Y*_96×*N*_ were sampled from a multinomial distribution for each sample with probability of *π*_,*n*_.

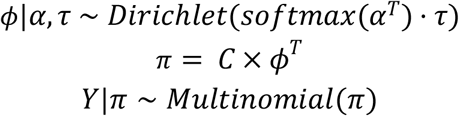

In the model, the standard deviation of *β* and the precision parameter *τ* are key prior parameters. *σ*_*β*_ was fixed as 0.5 as was decided from our observation of TCGA breast cancer data where the estimated posterior effect sizes of association of scaled covariates with signatures tended to be of standard deviation of 0.5. For *τ*, we assume a prior of lognormal distribution with log mean 0 and log standard deviation 2.

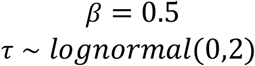

All analysis for *Diffsig* was performed in R v4.1.0 with a Bayesian hierarchical model specified in the Stan probabilistic programming language and the posterior inference was performed using the rstan package v2.21.3 with 4 cores and 4 chains^18^.

### Simulations

Simulated datasets were generated with number of donors (*N*) of 50, 100, 150, 200, and 1,000. In order to show that the model works with any type of risk factor, we generated in total 300 simulations: 20 simulations for three different types of risk factors (binary, categorical, and continuous) and for five different sample sizes (N=50, 100, 150, 200, and 1000). In order to evaluate if the model works in various contexts, we tested on mutational signatures known to be present in i) breast cancer (COSMIC SBS1, 2, 3, 5, 13) and ii) liver cancer (SBS 1, 4, 5, 12, and 29) found from previous studies^17^. Signatures were selected based on metrics from Alexandrov et al. (2020): proportion of each signature for each cancer sample from ICGA and TCGA datasets collected in the PCAWG consortium and the median mutation burden of each signature in samples that show that signature. Risk factors (*X*) in continuous and binary format were generated, sampling from either a normal or binomial distribution, respectively. To evaluate associations with categorical risk factors, a 3-level categorical variable was generated, sampling from a multinomial distribution, which then was modeled with an intercept and two dummy variables for the non-reference levels. We set the over-dispersion hyperparameter *τ* = 100, sampled associations *β* from a uniform distribution spanning [-2, 2], and centered the *β* values for the intercept and two non-reference level coefficients. The variance of the true associations was 4/3. The number of total mutations per donor, *J*, was sampled from a negative binomial distribution with mean 300 and dispersion parameter 1/100 in order to generate a typical mutation range as seen in whole-exome sequencing-based data from TCGA breast cancer dataset (e.g., within 10-500, excluding hyper-mutated samples). The observed 96 dimension mutation counts, *Y*, were then sampled from the model with fixed *C, X, τ* and sampled values for *β* and *J*. In total 20 simulations were generated, where for each iteration a new set of *β, X, J* were simulated.

To validate that our model would produce accurate estimates on all types of risk factors, we tested on binary, categorical, and continuous risk factors. In addition, analogous to linear or logistic regression, excluding a risk factor that is associated with the presence of one or more signatures may induce bias in estimates which can reduce the power of the test and/or lead to false positives. To empirically verify this possibility of bias from excluded covariates, we generated true values of (*α, β, ϕ, π*), assuming that there were two continuous risk factors associated with the signatures, where the risk factors were positively correlated, as they were sampled from a multivariate normal distribution (*β*∼*N*(0, Σ_2×2_), *where σ*_12_ = *σ*_21_ = 0·7). Two scenarios were tested: i) including the two continuous risk factors in *X* and ii) only one of the two risk factors in *X*.

After fitting the model using *rstan*, the convergence within and between 4 Markov chain estimates were examined with a standard MCMC diagnostic measure, the Gelman-Rubin statistic 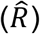 value, for each parameter^19^. The model chains were considered to be well converged when 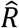 values were below 1.05. Posterior means of the parameters 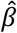 were compared to the true parameters *β*, and coverage and width of 80% credible intervals were assessed.

### TCGA Breast Cancer

TCGA mutation calls were retrieved from NCI GDC website in a MC3 file in hg19 coordinates and were then subset to 1,201 breast tumor samples to evaluate our model on real data. We selected single nucleotide variants (SNVs). We omitted germline chromosomes and adjacent normals. We then used the R package *VRanges* to generate the mutation count matrix from the MAF file^20^. Hypermutated samples with mutation counts ranging from 563 to 5037 were excluded from the dataset leaving only samples with mutation counts in the range of approximately 10-500 mutations. Breast cancer samples are commonly assigned to one of the five molecular intrinsic subtypes (PAM50) which are originally identified by clustering 50-gene expressions: luminal A (LumA), luminal B (LumB), HER2-enriched, basal-like, and normal-like^21^. In our analysis, we removed normal-like subtypes and only kept the remaining four.

Before applying *Diffsig* on the TCGA breast cancer dataset, the data was evaluated to determine whether varying sequence coverage and mutational burden induce heterogeneous uncertainty on estimated contributions for mutational signatures. This was done by selecting one of the HER2-enriched samples with sample ID TCGA-3C-AALI-01, which has a total of 428 somatic mutation counts. This mutation count was down-sampled using binomial sampling with a success probability of 0.1. We generated 100 different down-sampled mutation counts and estimated contributions on breast cancer COSMIC signatures using the non-negative least squares (NNLS) model from the *MutationalPatterns* R package^10^. Variance of the estimated contributions was assessed to see whether heterogeneous uncertainty exists.

We then applied *Diffsig* on TCGA samples to evaluate if the model would recover well-characterized associations. Along with the mutation counts, we obtained two risk factors for evaluation. One risk factor is the homologous recombination deficiency (HRD) scores, a metric that combines 3 different scores: HRD-LOH, LST (large-scale state transitions), and NtAI (number of telomeric allelic imbalances) scores, where HRD score > 42 is defined as HR deficiency ^22 23^. Another risk factor we considered was the PAM50 molecular subtype for 915 TCGA breast samples (basal-like n=161, HER2-enriched n=74, luminal A n=487, luminal B n=193)^24^. Along with the categorical variable coding for molecular subtypes, we tested on a continuous subtype variable derived from the correlation of the samples to the PAM50 subtype centroids to see how the results compared to analysis using the categorical subtypes^25^.

Risk factor variables were preprocessed based on the variable type as mentioned previously. However, for ease of comparing continuous to binary covariates, we did not scale the continuous HER2 subtype variable as it already had similar variance to the binary version. The mutation count matrix was subset to the 907 samples that have HRD scores and PAM50 subtype information available. Similar to the simulated datasets, 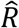 values of each parameter were examined to determine MCMC convergence.

### Model Evaluation

For simulations, we considered the model to accurately capture the true “*β*s” when i) the 80% credible intervals covered the true *β* at their expected rate and ii) the root mean squared error (RMSE) of the estimated *β*s was small.

For real data, the true *β* are not known, so instead, we evaluated the model qualitatively. The model was tested on two risk factors with previous knowledge to be associated with certain signatures. First, since a higher HRD score indicates homologous recombination deficiency, we tested the HRD score and COSMIC signatures, expecting a positive association between HRD score and SBS3 which is known to be associated with HRD. Second, Pitt et al. (2018) observed that tumors categorized as basal-like subtypes have significantly fewer APOBEC mutations compared to HER2-enriched subtypes. Therefore, we inspected whether the estimated associations show that HER2-enriched tumors had higher association with APOBEC signatures in COSMIC (SBS2,13) compared to basal-like tumors.

### Comparison with *HiLDA*

Unlike *Diffsig, HiLDA* estimates *de novo* signatures, which have a different format from the COSMIC signatures used in our analysis. Rather than summarizing in 96-dimension vectors, *HiLDA* estimates a signature with 3 sets of probabilities: probabilities of bases (A,C,G,T) in the flanking bases, and probabilities of mutations (C>A,C>G,C>T,T>A,T>C, T>G) (Supplementary Fig3). Therefore, *HiLDA* signatures were qualitatively compared to COSMIC signatures by comparing the flanking bases and mutations of *HiLDA* signatures to those of COSMIC signatures. In addition, *HiLDA* includes two different tests: the global test and the local test. The global test assesses whether there is a difference in the mean contribution between two groups with a Bayes factor across any of the *HiLDA* signatures, and the local test shows whether there is a difference in the mean contribution between two groups for each *HiLDA* signature. Note the mean contribution, or what is referred to in the HiLDA method as *exposure*, is the relative contribution for each binary risk factor of one group. In our benchmark, we compared with local test results in order to compare the associations between the risk factor and each signature.

## Results

### Simulation

Simulation datasets were generated to validate that our model can accurately capture the associations between risk factors and signatures. In simulating data from the model, we found that point estimates tended to have low error, and credible intervals (CIs) obtained their nominal coverage, across a range of risk factor types and sample sizes. We have seen that the mean coverage of the 80% CIs within each simulation was near 1 (>0.99) for all three types of risk factors regardless of the sample size (*R* <1.05, Supplementary Fig). As the sample size increased, the mean width of the CIs and the average RMSE of the estimated *β*s decreased as expected with a properly calibrated model (Fig2A,B). Nonetheless, even with a sample size of 50, the median of the average RMSE was equal to or less than 0.1, showing that our model is capable of achieving estimates approximate to the true values even with small sample sizes. Sample size increase led to an increase in computation time where 50 samples required less than 5 minutes, 200 samples required around 25 minutes, and 1,000 samples required 5-6 hours with 4 cores.

**Figure 2.**
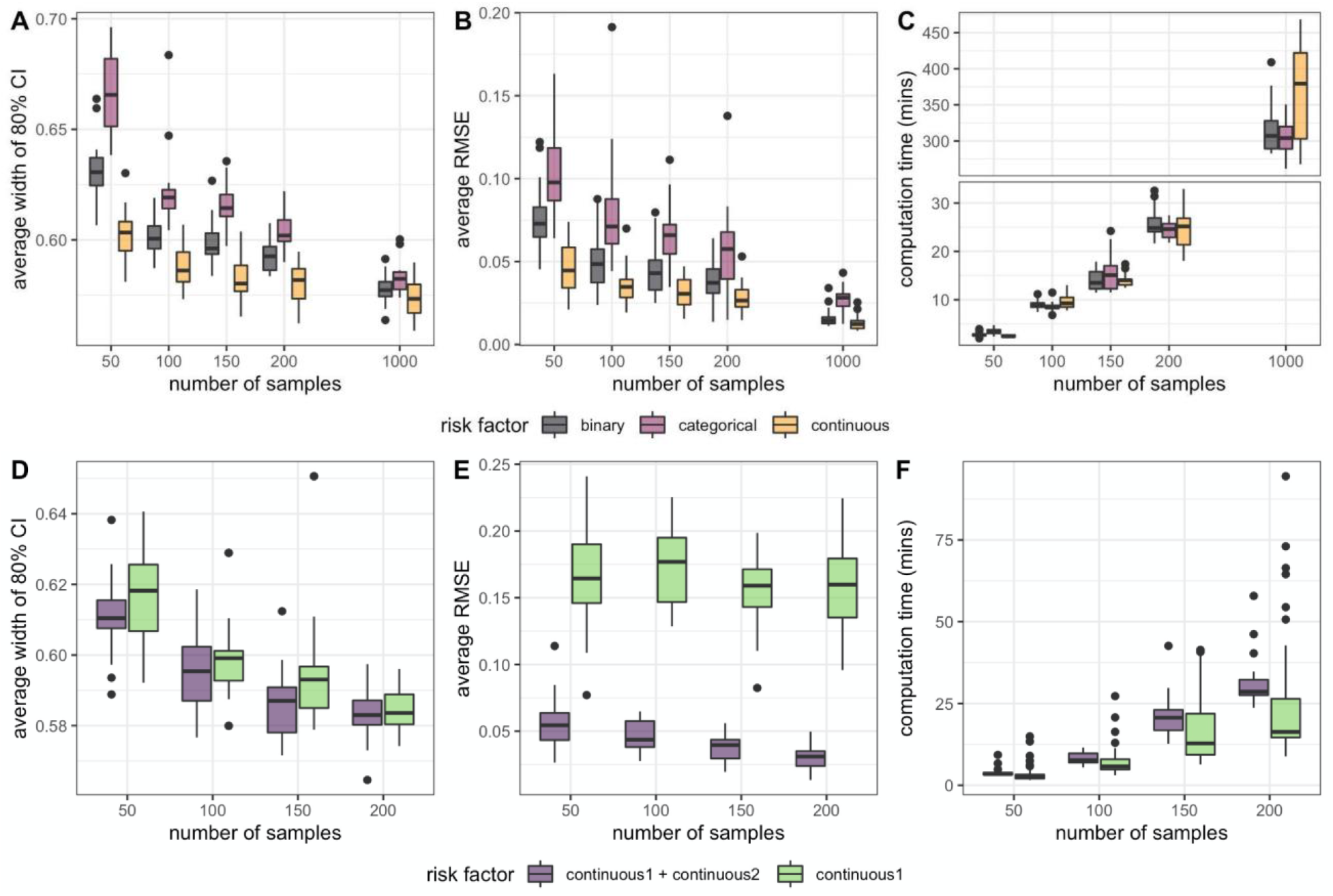
Simulation results across a range of risk factor types and sample sizes. A model including one risk factor (binary, categorical, or continuous) shows changes in (A) 80% credible interval coverage, (B) average root mean squared error (RMSE), (C) computation time (mins) as the number of samples increased. A model including all or only one of the risk factors shows changes in (D) 80% credible interval coverage, (E) average RMSE, (F) computation time (mins) as the number of samples increased.

Results showed that including both risk factors that influenced the presence of signatures had higher interval coverage (Supplementary Fig1C,D) and significantly lower average RMSE, compared to a model that only included one of the risk factors. This showed that including multiple risk factors that are believed to be jointly associated with the signatures can lead to more accurate estimation of the associations compared to modeling with only one risk factor. Moreover, as the number of samples increased, the average width of the 80% credible interval and average RMSE decreased (Fig2D, E). Within the same sample size, the average widths of simulations with two covariates are lower or similar to ones with only one covariate. Computation time was higher when including both risk factors compared to only including one risk factor (Fig2F). However, considering the increase in the number of parameters, there was only a moderate amount of increase in computational cost.

In addition, we tested *Diffsig* on simulations with another set of mutational signatures associated with liver cancer^17^. Results were similar to what was observed with breast cancer signatures; again, including all potential risk factors was less biased than only including one of them (Supplementary Fig2).

### TCGA Breast Cancer

*Diffsig* has been developed to avoid the inherent sampling variability, instead allowing association tests with various types of risk factors using a hierarchical Bayesian model. Methods using point estimates for estimated contributions to binary dependent variables may underestimate the sampling variability. To empirically demonstrate such a drawback, we performed a naive analysis using NNLS on one of the basal-like TCGA breast cancer samples with five breast cancer-related signatures (SBS1, 2, 3, 5, and 13). It was observed that the relative contributions (signature X’s contribution/sum of all signatures’ contribution) varied across different down-sampled datasets (Fig 4A). For SBS13, the relative contribution ranged from nearly 0.2 to over 0.65 for certain down-sampled datasets.

**Figure 4.**
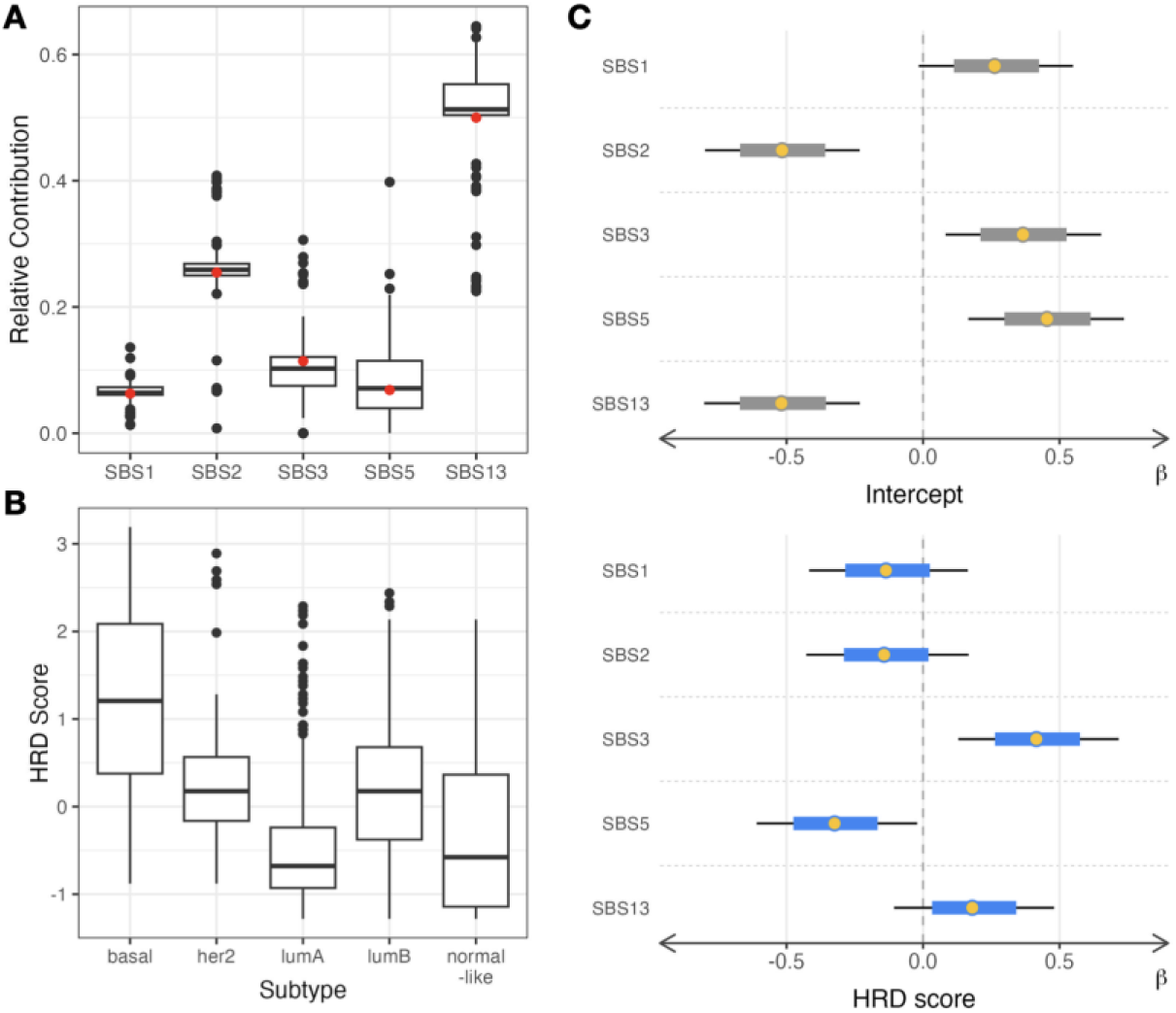
(A) Relative contribution from NNLS on down-sampled experiments of a basal sample from TCGA-BRCA dataset with breast cancer-related COSMIC signatures (red dots represent relative contributions generated with original mutation counts) (B) Homologous Recombination Deficiency (HRD) score distribution in breast cancer subtypes (C) Diffsig results including median (yellow dots), 80% and 95% credible intervals (blue and black lines) for the intercept (grey, top) and HRD score (blue, bottom).

Moreover, *Diffsig* was able to detect the associations across various risk factors which would be expected from previous literature on somatic mutations and breast cancer. From the preliminary exploratory analysis of the TCGA breast cancer data, tumors with basal-like subtypes have a higher average HRD score than the other subtypes (Fig4B). The basal-like subtype has previously been shown to have a high association with HRD mutation signature SBS3^17 26^. With this knowledge, we tested the association of HRD score with the five mutational signatures previously reported as common in breast cancer with *Diffsig* (Fig4C). The estimated association *β* for the HRD score was much higher in SBS3 than the other 4 signatures. Also, the high estimated *β* from the intercept of SBS5 is indicative that this signature was prevalent in most cancer patients^17^. All of the 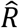 values for the estimates were below 1.05 (≈1.00) and the total computation time was around 12 hours for 931 samples.

To compare our performance with HiLDA, we tested both *Diffsig* and *HiLDA* on a binary risk factor (subtype). After selecting only the HER2-enriched and basal-like samples from TCGA, both HiLDA and Diffsig were run on a binary indicator of HER2-enriched samples. Although HiLDA does not produce 96-dimensional mutational signatures, we were able to qualitatively align the HiLDA signatures 1, 2, 3, and 5 with the COSMIC signatures 1, 2, 3, and 13 based on their dominant features (Fig5A, Supplementary Fig 3).

Once we ordered the *HiLDA* signatures to match the order of the corresponding COSMIC signatures, we observed that the estimated difference in mean exposures (or contribution) from *HiLDA* aligned with the ranks and directions of the estimated associations from *Diffsig* (Fig5B, C). Note that the scales of the x-axis differ as the models are inherently different in that *Diffsig* measures associations while *HiLDA* measures differences in contributions. HER2-enriched samples, when compared to basal-like samples, had a higher association with SBS2 and SBS13 than with other signatures. This agrees with our previous knowledge that HER2-enriched samples tend to have higher APOBEC mutagenesis compared to basal-like samples^26^. Both results showed convergence 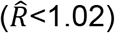, while the computation time was slightly faster with *Diffsig* compared to *HiLDA* (*Diffsig* 58 mins, *HiLDA* 78 mins with 241 samples).

Aside from considering subtypes as a categorical variable, we considered continuous measures of subtypes using the correlation to subtype centroids^25^. Estimated associations when considering the continuous correlation to HER2-enriched subtype were almost identical to the estimated associations for binary HER2-enriched subtype status (Figure 5C,D). Estimates using the binary definition of subtype had reduced computation cost (6.65 mins, 242 samples). In addition, the same process was tested on another set of breast cancer-related COSMIC signatures (SBS2,3,5,8,13) to assess whether the selected set of signatures matter (Supplementary Fig4). Results show similar results as to having high associations in HRD score with SBS3 and HER2-enriched samples with SBS2 and 13.

**Figure5.**
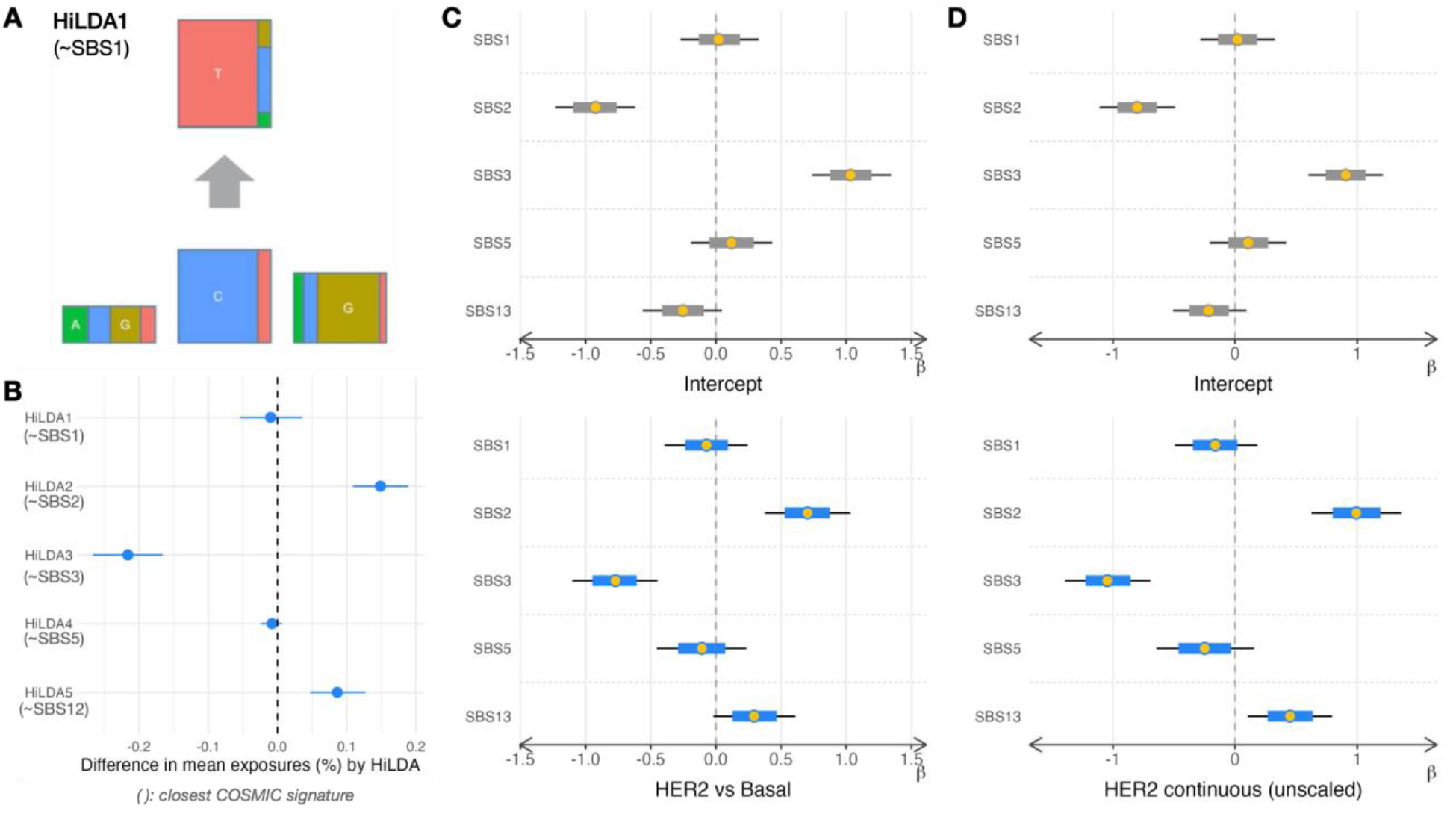
*Diffsig* and *HiLDA* results on a subset of HER2-enriched and Basal-like subtypes from TCGA-BRCA (A) Each *HiLDA* signature was connected to a COSMIC signature, e.g. *HiLDA1* has similar composition of SBS1 (B) *HiLDA* results with error bars representing the difference in mean exposures (in percentage) for each *HiLDA* signature (C) *Diffsig* results including median estimates (yellow dots), 80% credible intervals (grey/blue lines), and 95% credible intervals (black lines) for estimated associations between binary HER2-enriched subtype indicator and COSMIC breast cancer signatures (SBS1,2,3,5,13) (D) *Diffsig* results for estimated associations between continuous HER2-enriched subtype correlation and COSMIC breast cancer signatures.

## Discussion

A key task in cancer genomics is to identify the etiology of discovered mutational signatures by relating them to risk factor data. *Diffsig* allows researchers to quantify and perform inference on the associations of diverse risk factors with reference mutational signatures, leading to better understanding of the tumor development process and improved models of tumorigenesis. From simulation, we observed that when the sample size was sufficient, i.e. more than 50 samples given the simulation parameters, our model was able to correctly capture the estimates within the 80% credible intervals and with sufficiently low RMSEs. Real data results from TCGA support that our model is capable of estimating expected associations in a variety of contexts. Note that such results with high power require sufficient sample sizes along with a balanced design if testing on discrete covariates.

*Diffsig* is also capable of estimating the associations in real data applications, as shown with the TCGA breast cancer dataset. These results aligned with what has been seen in previous studies of breast cancer related to HRD and APOBEC activity. Homologous recombination deficiency in general are assessed based on various indexes, e.g. mutational signatures, DNA-based measures of genomic instability ^27^. In previous breast cancer studies, it was shown that patients with high contribution of COSMIC signature 3 (SBS3) as well as those with high HRD scores were considered homologous recombination deficient which may indicate high sensitivity to certain treatments, e.g. PARP inhibitor^3,28 29^. *Diffsig* showed consistent associations between these two measures. In addition, a study by Roberts et al.(2013) was performed on 14 cancer types including breast cancer with whole-genome and whole-exome sequencing datasets including TCGA^30^. This study showed that HER2-enriched samples have significantly higher contributions of APOBEC-related mutations compared to other PAM50 subtypes. The contrast was most clearly seen between HER2-enriched and Basal-like subtypes, also corresponding to what we observed in our analysis. Compared to previous methods which have emphasized mutational signature discovery, *Diffsig* is focused on evaluating established signatures. *Diffsig* does not involve a *de novo* signature identification step, but instead leverages the well-established mutational signatures from existing catalogs, e.g. COSMIC. To the extent that current catalogs contain the relevant signatures for the samples under examination, leveraging these catalogs facilitates the comparison of associations across studies. However, for data sets with sufficiently large number of samples, NMF can be trained on the dataset instead of using existing reference signatures. Although we have limited our scope here to single-basepair substitutions (SBS), *Diffsig* is capable of taking other categories of mutations such as double-based substitutions (DBS) or inserts and deletions (indels).

*Diffsig* is operationally well-aligned with current understanding of mutational signatures. Specifically, *Diffsig* uses a Dirichlet-Multinomial distribution for the observed counts following *HiLDA*, which reflects our belief that the mutations in a sample are a composite of multiple underlying mutational signatures. The Dirichlet layer of the model allows for overdispersion of the sample-signature contributions, such that two samples with the same risk factor values may have different latent contributions ɸ. One layer of our *Diffsig* model maps from the sample-signature associations *α* = X*β* to the sample-signature contributions ɸ. To accomplish this, a softmax function is applied which takes input in the domain of R^n^ and outputs values on the n-simplex, an (n-1) dimensional subspace of [0,1]^n^. One property of the softmax function is that any constant shift applied to all elements of *α* results in the same output. Therefore, it is necessary to apply a zero-centered prior to *β* to provide identifiability. Additionally, ɸ will exhibit the properties of other compositional data, where an increase in the contribution of one feature (here signature) necessitates decreases in the other features. Therefore, we recommend a compositional interpretation of estimated *β*s within each risk factor, while per-feature elements of *β* should not be compared across risk factors. In other words, we may conclude that one signature is more or less associated with a risk factor than other signatures, but we cannot conclude that a signature is more associated with one risk factor than the other risk factors. Another restriction we set is the sampling distribution of *β*. We learned from TCGA dataset that the posterior *β* converged to values within the standard deviation of 0.5 hence fixed *β* to be modeled as a zero-centered normal with a fixed standard deviation of 0.5 to regularize the parameter to reasonable values. Adding further layers to the hierarchical model with regard to the variance of *β* is possible but we found led to much longer convergence times during MCMC sampling.

For our real data analysis, we have limited our analysis to clinical tumor samples and did not incorporate matched normal samples. By including healthy tissue samples from the tumor patients, our analysis will enable us to determine somatic mutations from germline mutations which would allow more accurate identification of risk factors at the population level by adjusting patient-specific mutational patterns.

## Acknowledgments

The authors would like to acknowledge the following individuals for helpful comments and suggestions on the work: Zhi Yang and Jean Fan. This work was supported by the NIEHS grant P30-ES010126 and NCI grant R01-CA253450.

## Data availability

The results published here are in whole or part based upon data generated by The Cancer Genome Atlas Research Network: https://www.cancer.gov/tcga. The MC3 MAF file used to generate the somatic mutation count data in this paper can be found at the GDC Application Programming Interface (API): http://api.gdc.cancer.gov/data/1c8cfe5f-e52d-41ba-94da-f15ea1337efc.

*Diffsig* is publicly available as an R package (https://github.com/jennprk/diffsig).

