## Supplementary materials for "Diffsig: Associating Risk Factors With Mutational Signatures"

**Supplement 1: Simulation Results 80% Credible Interval Coverages**

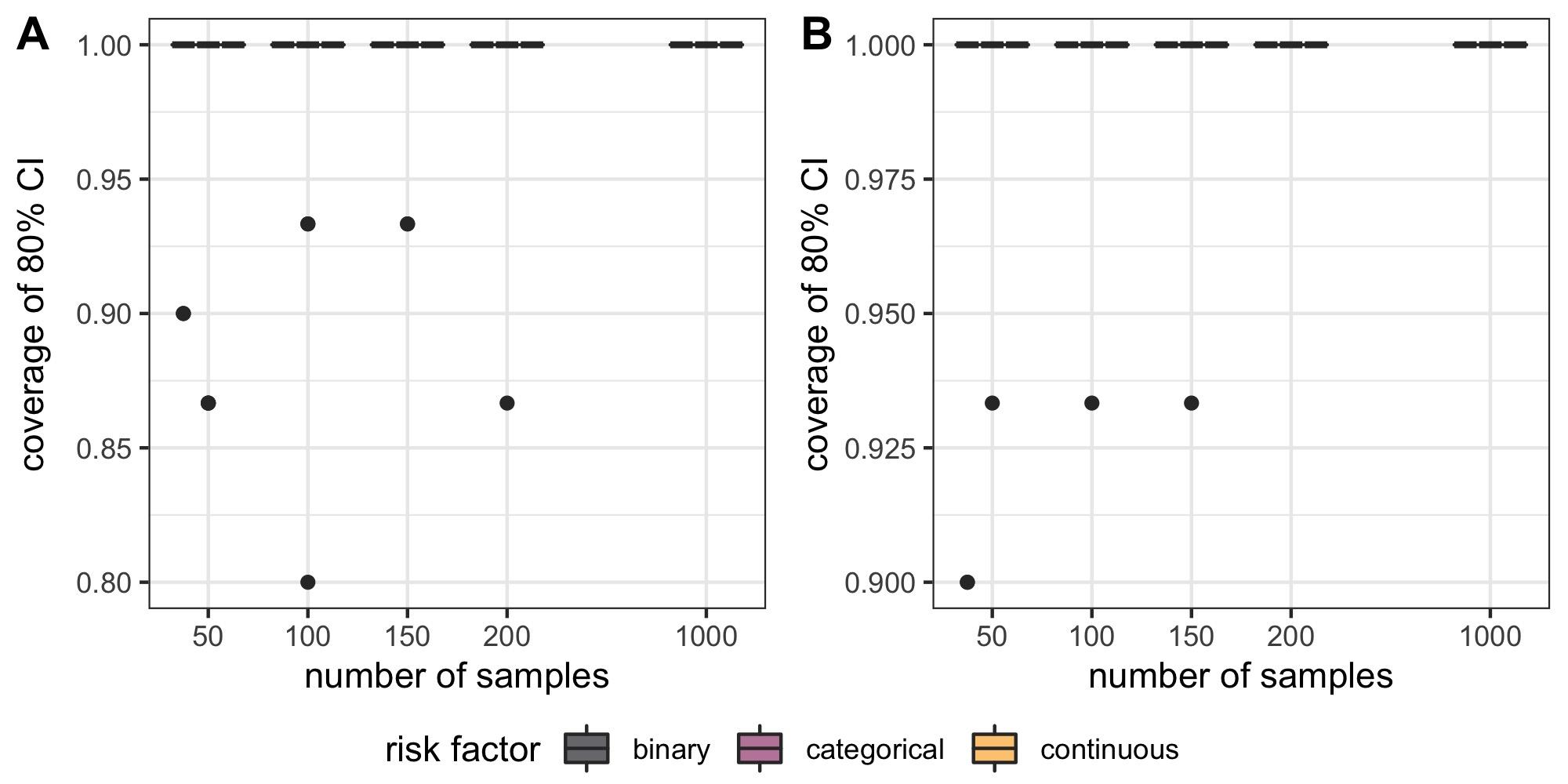

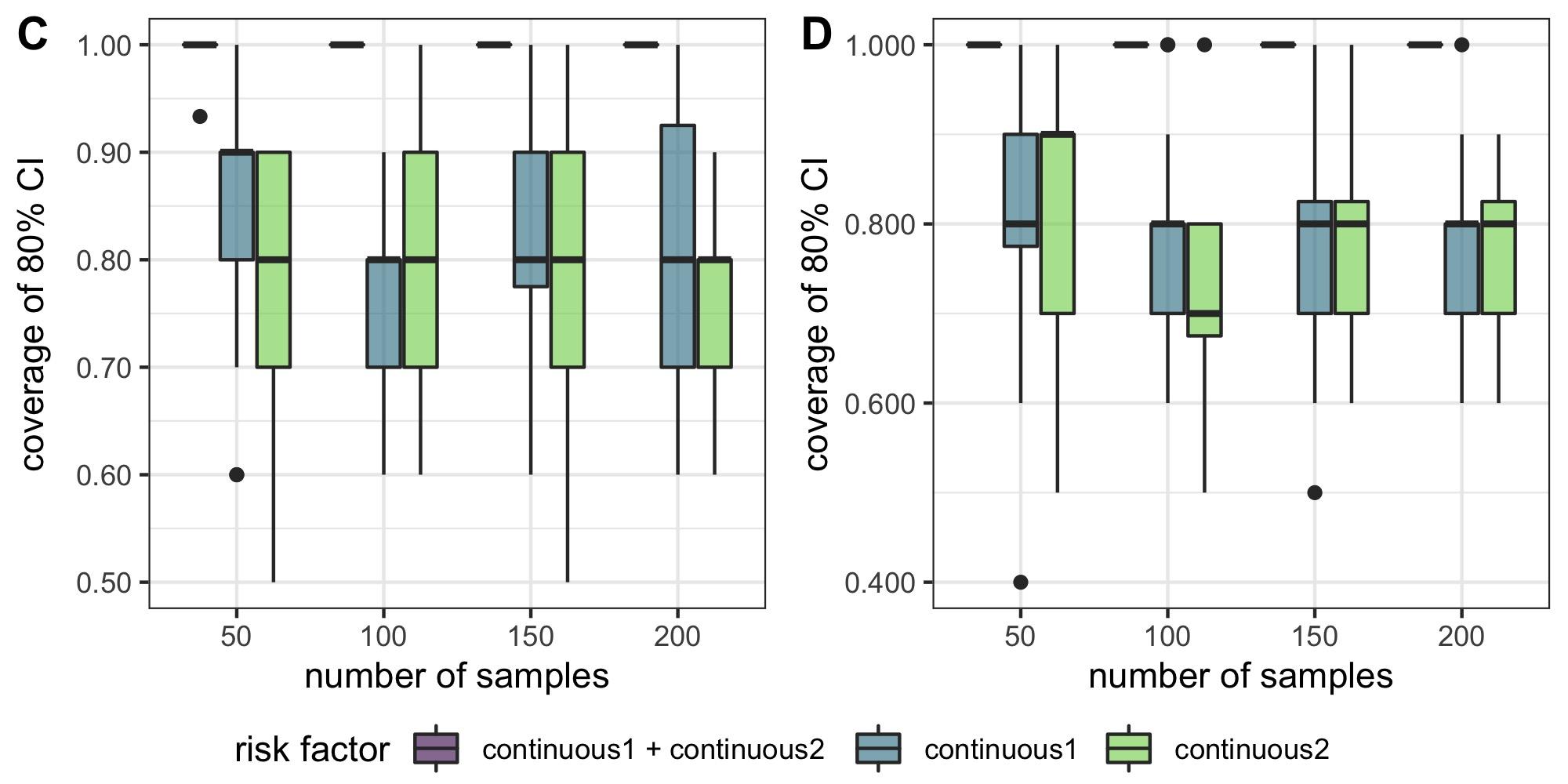

Most of the simulations based on (A) breast cancer signatures and (B) liver cancer signatures were able to capture the true $\beta$ within the 80% credible interval. On a scale of the coverage 0 to 1, most of the univariate risk factor models were able to estimate all $\beta$ in the 80% CI (coverage=1). Even for simulations that were unable to fully capture the true $\beta$, the coverage was greater or equal to 80%. This also improved as the number of samples increased in both signature sets.

When comparing the coverage of two risk factors to either one of the risk factors, we could see that including both truly associated risk factors have higher coverage of 80% CIs. Coverages of the first case are approximately 1 while including either risk factor yielded a median of 0.9 or 0.8 for (C) breast cancer signature-based simulations, and the coverage even went down to a median of 0.7 in cases of (D) liver cancer signature simulations. This result provides supporting evidence that when more than one risk factor is expected to be associated with cancer or cancer-related mutational signatures, it is encouraged to include all instead of running univariate analyses.

**Supplement 2: Simulation Results with Liver Cancer Signatures**

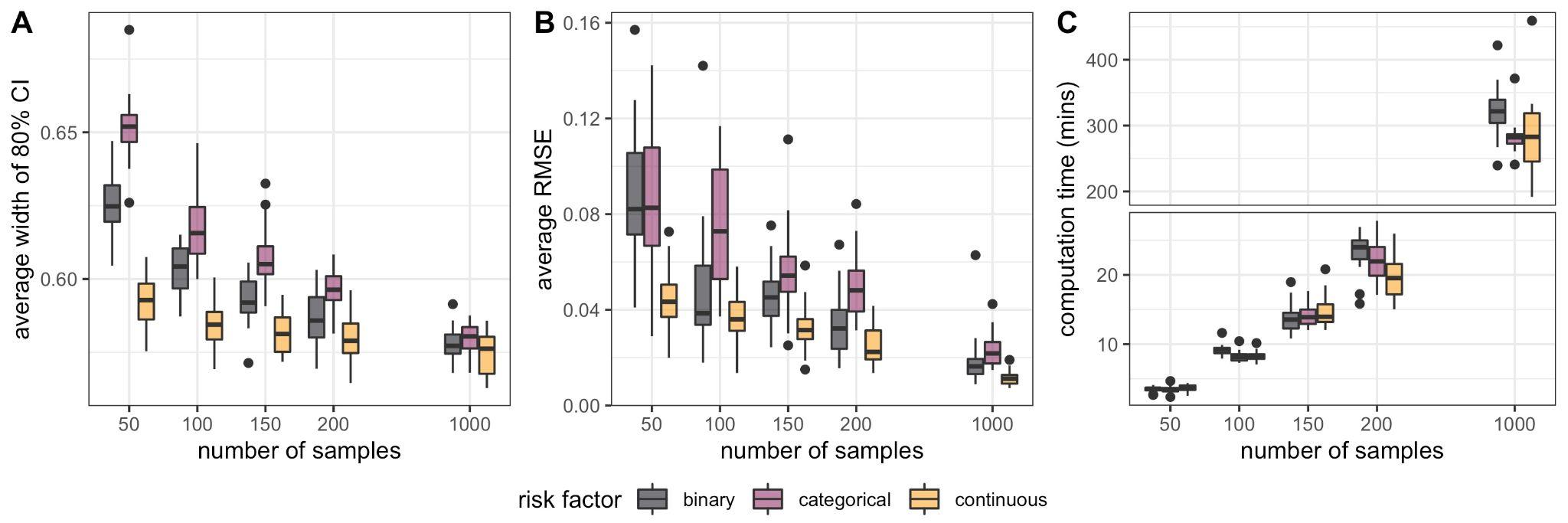

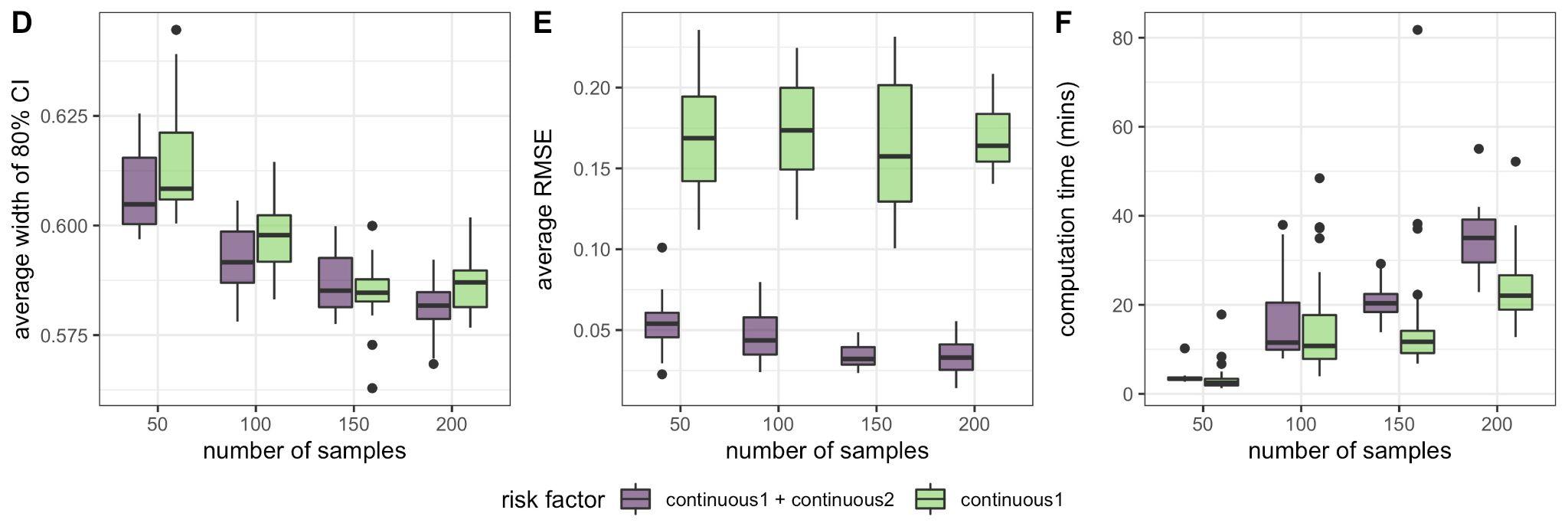

In order to ensure our model works on other subsets of signatures, we generated the simulation results using liver cancer-related signatures (SBS 1, 4, 5, 12, and 29).

Results showed similar results where the mean coverage of the 80% CIs within each simulation was near 1 (>0.99) for all three types of risk factors regardless of the sample size. The mean width of the CIs decreased as the sample size increased which shows a more precise estimation is capable with a higher number of samples (A). In addition, as the sample size increased, the average RMSE of the estimated $\beta$ dropped while computational cost increased (B,C).

Moreover, multiple risk factors on liver cancer signatures with *Diffsig* also showed more accurate estimation of the associations compared to modeling with only one risk factor (D,E) along with only moderate amount of increase in computational cost (F).

**Supplement 3 - *HiLDA* Signatures on TCGA Breast Cancer Data**

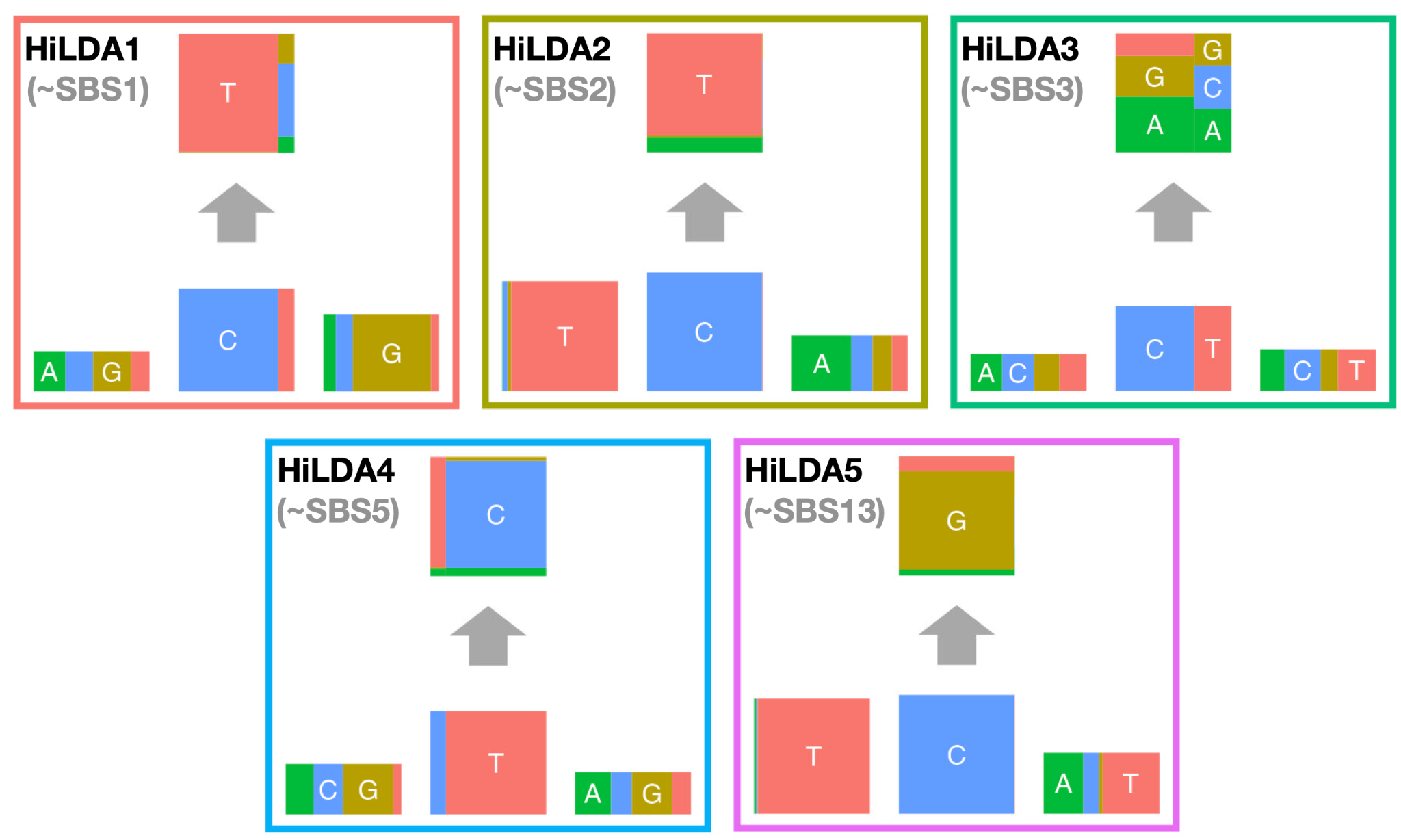

Above are the five latent signatures that were generated from HiLDA based on the TCGA breast cancer dataset. When compared to the COSMIC signatures, HiLDA signature1 highly resembles COSMIC SBS1 in which C>T mutations are dominant and with high guanine proportion for the following base. HiLDA signature2 resembles SBS2 which has 99% C>T mutations while having 98% thymine in the preceding base. HiLDA signature5 also appears to match well with SBS13 where 99% CpN mutations in which around 80% belongs to C>G mutations while preceding base is 94% thymine and following base is 44% adenine and 43% thymine. In addition, HiLDA signature3 is somewhat similar to COSMIC SBS3 where mutations occur fairly across all 96 mutation contexts. *HiLDA* signature4 could be considered to correspond to SBS5 as T>C and C>T are the major mutations, however, it does not have a clear one-to-one connection with a COSMIC signature unlike the other ones.

**Supplement 4 - TCGA results with a different set of breast cancer signatures**

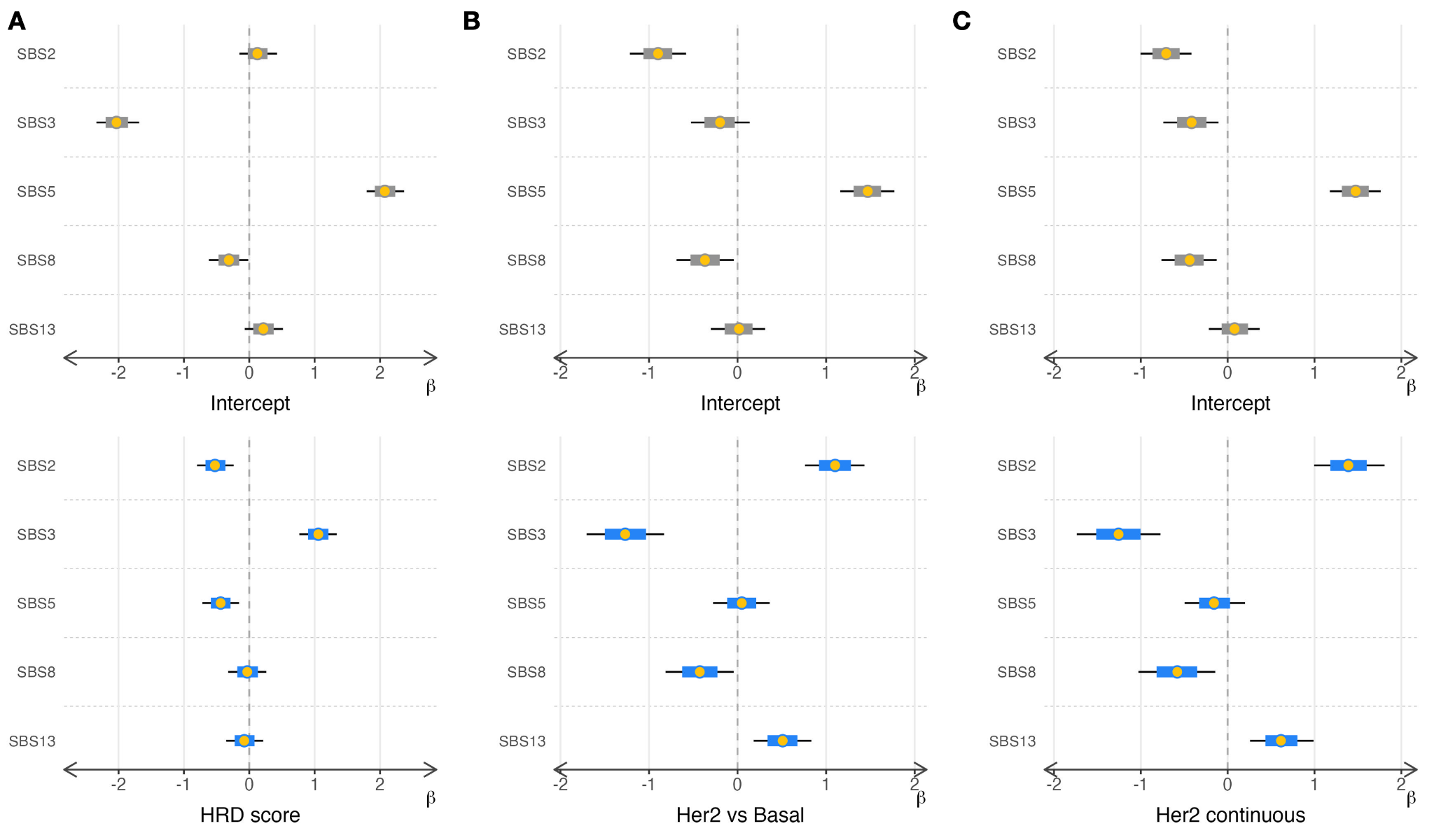

Here, we excluded SBS1 and replaced it with SBS8 to see if *Diffsig* is capable to detect the expected association even with different signatures. It shows that HRD score still has the highest association with SBS3 relative to other signatures (A), and both binary HER2 subtype indicator and continuous HER2 subtype risk factors have high associations with APOBEC-related signatures; SBS2 and 13.
